# Gait analysis of healthy rhesus monkeys using a pressure-sensing walkway system

**DOI:** 10.1101/2020.04.14.040907

**Authors:** Jinyoung Won, Keonwoo Kim, Hyeon-Gu Yeo, Junghyung Park, Jincheol Seo, Yu Gyeong Kim, Sung-Hyun Park, Won Seok Choi, Chang-Yeop Jeon, Kyung Seob Lim, Jae-Won Huh, Young-Hyun Kim, Jinwoo Jeong, Ji-Woong Choi, Youngjeon Lee

## Abstract

Gait analysis in non-human primate models has been performed to elucidate the neural systems involved in controlling quadrupedal locomotor behavior. This study aimed to use a pressure-sensing walkway to identify characteristics of species-specific quadrupedal locomotion in rhesus monkeys. A total of nine healthy adult female rhesus monkeys *(Macaca mulatta)* were used for gait analysis. We measured the temporo-spatial and kinetic parameters of quadrupedal gait using a custom-built pressure-sensing walkway and compared the left- and right-side parameters to assess the symmetry of the gait pattern. All temporo-spatial and kinetic parameter values showed no significant differences among the nine rhesus monkeys for both the forelimbs and hindlimbs. However, significant differences were observed between forelimb and hindlimb kinetic parameters such as peak vertical force, vertical impulse, and the percentage of body weight distribution. All kinetic parameter values were higher for the forelimbs than for the hindlimbs. These data indicated that the center of gravity in healthy rhesus monkeys is located at the forelimbs rather than at the hindlimbs while walking. Furthermore, the symmetry indices considered for symmetric gait pattern showed a low variability. Most median symmetry index values were nearly zero, indicating no difference between the right and left sides. This study described valid methods for assessing gait parameters and demonstrated rhesus-specific characteristics of quadrupedal locomotion, providing a basis for the assessment of gait normality in rhesus monkeys.

## Introduction

Walking is one of the most common physical activities and plays an important role in daily life [1]. Gait analysis has been widely used to assess an individual’s walking ability and identify posture and movement problems. The main causes of gait disorders include musculoskeletal problems (e.g., diseases of joints, bones, or tissue) and neurological diseases (e.g., impairments of the sensory or the motor system), which directly affect daily activities [2, 3]. Differential diagnosis based on musculoskeletal and neurological examinations is therefore required to treat gait disorders using appropriate and effective therapy.

Abnormal or pathological locomotor activities can also be determined by functional gait assessment. As gait disorders are commonly associated with dysfunction in motor control and linked to sensory impairment, characterization of gait pattern can be useful in assessing functional recovery. In addition, gait analysis can be translated easily from laboratory animals to humans. Rhesus monkeys are genetically close to humans, and the general structure of their neuroanatomical system is more similar to that of humans than rodent systems are [4, 5]. Therefore, rhesus models of spinal cord injury, stroke, and neurodegenerative diseases have been used to investigate pathological mechanisms and assess therapeutic treatments in preclinical studies.

Gait analysis in non-human primates with habitual quadrupedal and bipedal locomotion has also been employed for the assessment of neurological deficits [6–8]. Grabli et al. used posture parameters to identify pathological gait patterns in Parkinsonian monkeys while the animals walked down a hallway [9]. Capogrosso et al. used a kinematic analysis system to analyze walking disorders and balance deficits that occurred after spinal cord injury in rhesus monkeys [10]. Nevertheless, although various assessment devices and gait parameters have been reported for the evaluation of gait function [11–14], temporo-spatial and kinetic walking characteristics of rhesus monkeys have not been investigated. In the present study, we described the walking gait of healthy rhesus monkeys and developed a normal gait database using a pressure distribution sensor apparatus.

The pressure-sensing walkway system is a pressure distribution sensor device that has been used to measure diverse gait values, including temporo-spatial and kinetic parameters. A symmetry index (SI) of each parameter considered for gait balance can be used as an indicator of gait normality. Recent studies have suggested that animal gait analysis using the pressure-sensing walkway system is less time consuming and is especially useful for repeated gait assessments [15]. In addition, previous animal gait studies have utilized the pressure-sensing walkway system and validated the methodology for use with multiple species such as horses, goats, sheep, dogs, cats, and Sprague Dawley rats [16–18]. However, such sensor systems have not been validated for use with the rhesus monkey, and the corresponding gait measurements have not been investigated. The primary aim of this study was to validate the use of a pressure-sensing walkway system that enables automated calculation of an array of gait parameters. The secondary goal was to evaluate the temporo-spatial and kinetic parameters for gait analysis and for assessing quadrupedal locomotion in clinically healthy female rhesus monkeys. We hypothesized that the temporo-spatial and kinetic parameter gait data would show no difference among the individual animals, and the SI of these parameters would have a low variability, indicating symmetric gait patterns of healthy rhesus monkeys. The results of this preliminary study could potentially help to differentiate between normal and abnormal gait.

## Methods

### Animals

We used nine adult (8-year-old) rhesus macaques *(Macaca mulatta)* obtained from Suzhou Xishan Zhongke Laboratory Animal Co. (Suzhou, China) and housed in individual indoor cages at the National Primate Research Center of the Korea Research Institute of Bioscience and Biotechnology (KRIBB) as described previously [19, 20]. The individual characteristics of the animals, including animal ID, age, gender, and body weight, can be found in S1 Table. Animals were fed commercially available monkey feed (Harlan, USA) supplemented with various fruits and water *ad libitum.* The controlled environmental conditions were as follows: temperature, 24 ± 2° C; relative humidity, 50 ± 5%; and a 12 h light/dark cycle. All procedures were approved by the KRIBB Institutional Animal Care and Use Committee (KRIBB-AEC-19046), and all experiments were performed in accordance with the relevant guidelines and regulations. All animals were monitored daily and were provided appropriate veterinary care by trained personnel.

### Experimental design

The experimental diagram of the gait assessment system is depicted in Fig 1A. Briefly, the testing apparatus for functional gait assessment consisted of a custom-built transparent plexiglass tunnel (3.21 m × 0.50 m × 0.79 m), a camera recording video images, and a pressure-sensing walkway for tracking the pressure applied by the four limbs of the monkeys.

**Fig 1.**
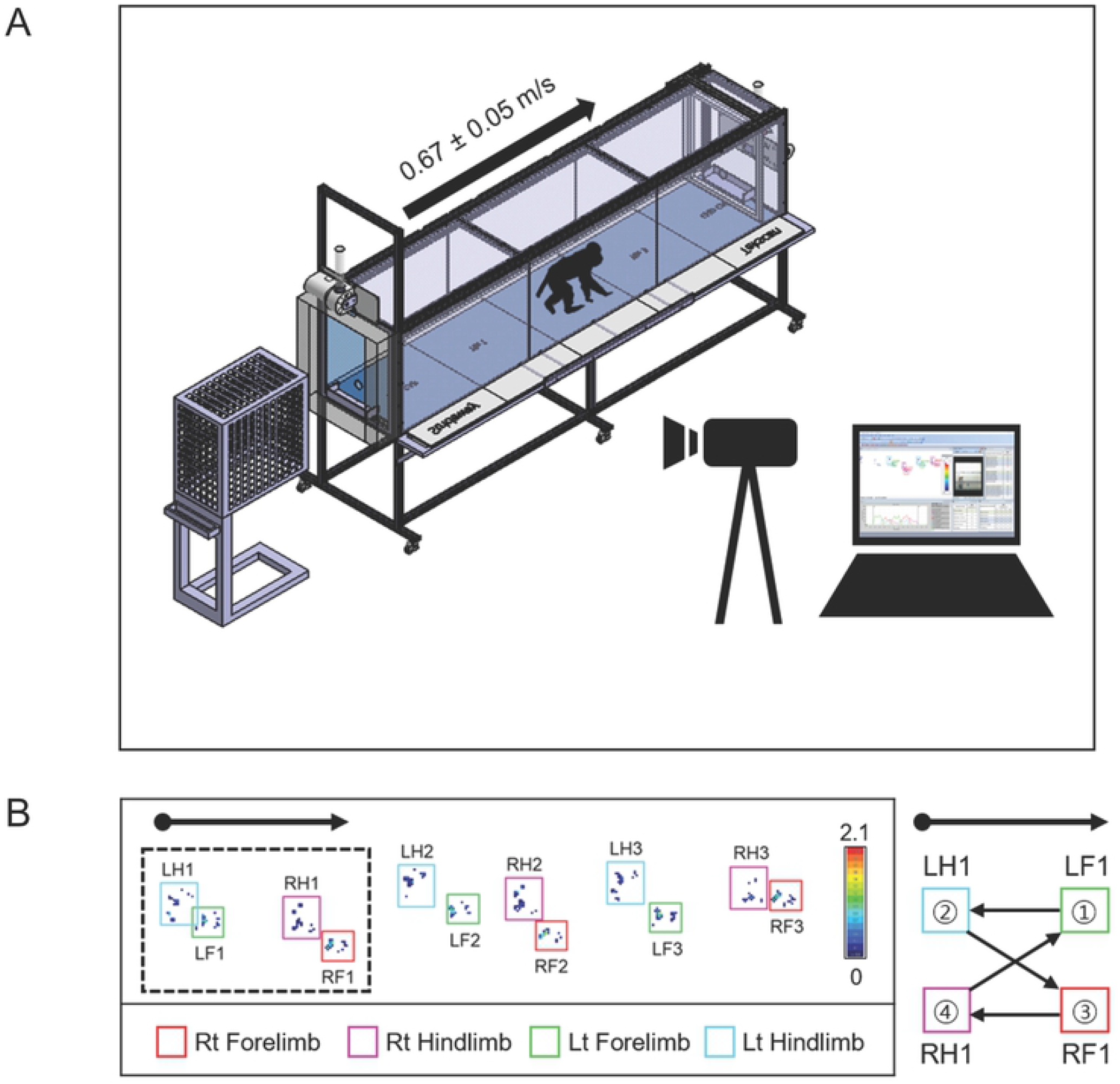
Experimental setup and gait analysis. (A) Schematic diagram of the gait assessment system consisting of a custom-built tunnel, pressure sensor mat, digital video camera, and dedicated software program. (B) Placement of each limb was detected by the sensor mat. The arrow indicates the direction of movement. The symmetrical gait of the rhesus monkey described in the right panel was presented when the ipsilateral hindlimb followed the leading forelimb.

### Functional gait assessment protocol

The monkeys were familiarized with the tunnel in order to reduce stress. As gait velocity is an important factor for the interpretation of gait data, the animals were pre-trained to maintain a constant speed between 0.6 and 0.7 m/s. Gait analysis was performed once a week for 1 month. Before data collection, each monkey was weighted on the same electronic scale for equilibration and calibration of the sensor device. The animals were stimulated to walk in a straight line on the pressure-sensing walkway mat by using reward pellets (Fruit Crunchies, Bio Serv, USA). The overall process frame was recorded using a digital video camera (HDR-CX405, Sony Corp., Japan) fixed on a tripod in front of the gait assessment system. The temporo-spatial and kinetic data were measured using the Strideway system pressure sensor mat (3-Tile High Resolution Strideway System, HRSW3; Tekscan Inc., South Boston, Massachusetts, USA), and its dedicated software program (Walkway 7.7; Tekscan Inc., South Boston, Massachusetts, USA) was used for data acquisition and processing. The Strideway system positioned inside the tunnel consisted of an active sensing area (1.95 m × 0.65 m) and end caps aligned with both ends of the sensing area. The supplementary video of monkey gaits that pre-trained to walk on the pressure-sensing walkway can be found in S1 Video. Placement of each of the four limbs detected by the pressure sensor mat was automatically visualized using the Walkway software program (Fig 1B and S2 Video). The tracks were manually marked by the strike box used to differentiate between the left and right forelimbs and hindlimbs. Data from an average of 40 trials were obtained for each monkey. Subsequently, data from the first five valid trials were selected and analyzed. A trial was considered valid if the monkey performed habitual quadrupedal locomotion without turning its head or resorting to bipedal locomotion.

### Gait analysis parameters

Data for the temporo-spatial and kinetic gait parameters and SI were collected. The temporo-spatial parameters, including stance time, swing time, stride time, stride length, percentage of stance, and percentage of swing were determined for each limb. Stance time was defined as the period of contact of the paw with the mat. Swing time was the period during which the paw was not in contact with the mat. Stride time was the time interval between two consecutive contacts of the same limb with the mat. Percentage of stance was defined as stance time/gait cycle time × 100. The percentage of swing was defined as swing time/gait cycle time × 100. Kinetic parameters, such as the peak vertical force (PVF), vertical impulse (VI), and the percentage of body weight distribution (BWD), were also determined. The PVF and the VI were normalized to the individual monkey’s body weight and represented as percentages of body weight. The percentage of BWD among the four limbs was defined as PVF of the specific limb/total PVF of the four limbs × 100. The SI of these gait parameters between the right side and the left side in both forelimbs and hindlimbs was calculated as previously described [21] and defined as follows: 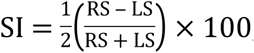, where RS represents the right-side parameter value and LS represents the left side parameter value. A negative value of SI indicated left side asymmetry. A positive value of SI indicated right side asymmetry. A near zero SI value indicated near complete gait symmetry.

### Statistical analysis

Statistical analyses were performed using SPSS 18.0 (SPSS Inc., Chicago, IL, USA). The data from nine rhesus monkeys (n = 9) were represented as the mean ± standard deviation. Coefficients of variation (CVs) were calculated to determine the average scattering. The Mann-Whitney test was conducted to evaluate the SI. Kruskal-Wallis test was used for comparing gait parameters of the left and right forelimbs and hindlimbs. A*p* value of < 0.05 was considered statistically significant.

## Results

### Temporo-spatial and kinetic parameter data of individual rhesus monkeys

Nine rhesus monkeys, judged to be clinically healthy based on the results of physical, orthopedic, and radiographic examinations, were enrolled and pre-trained to walk across the custom-built tunnel. A frame image extracted from a video recording shows a representative sequence of habitual quadrupedal locomotion (Fig 2A). The graph of force on the surface of the pressure sensor mat for each limb was obtained from the marked tracks (Fig 2B). Temporal parameters, including stride time, swing time, and stance time, are shown in Fig 2C. Gait data for the temporo-spatial and kinetic parameters of individual rhesus monkeys while walking over the pressure sensor mat can be found in Fig 3 and S1-S4 Tables. Individual data directly measured from each of the nine rhesus monkeys are presented as scatter plots in Fig 3. The average gait velocity of the monkeys on the pressure sensor mat was 0.67 ± 0.05 m/s during quadrupedal walking. Both the forelimb and hindlimb temporo-spatial data did not differ significantly among the monkeys (S2-S3 Tables). The forelimb and hindlimb kinetic data also did not differ significantly among the monkeys (S4-S5 Tables).

**Fig 2.**
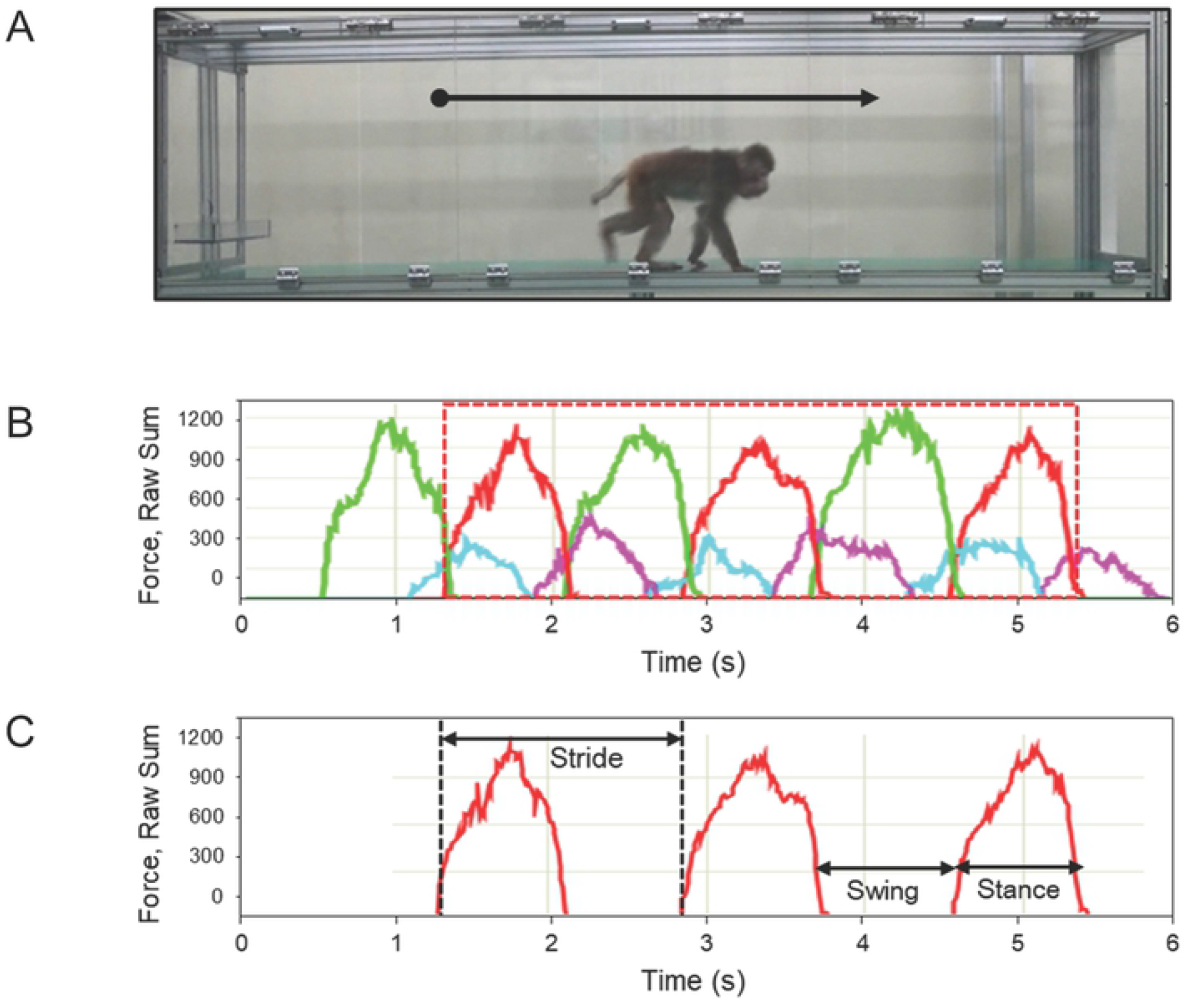
Representative recording of rhesus monkey. (A) Video image of a rhesus monkey walking on the pressure-sensing walkway. (B) Graph of force on the mat surface for each limb over time. (C) Right forelimb data obtained from the region outlined in red in (B) are presented. Stride time is the time interval between two consecutive contacts of the same limb. Swing time is the duration while the paw is not contact with the mat. Stance time is the contact period on the mat.

**Fig 3.**
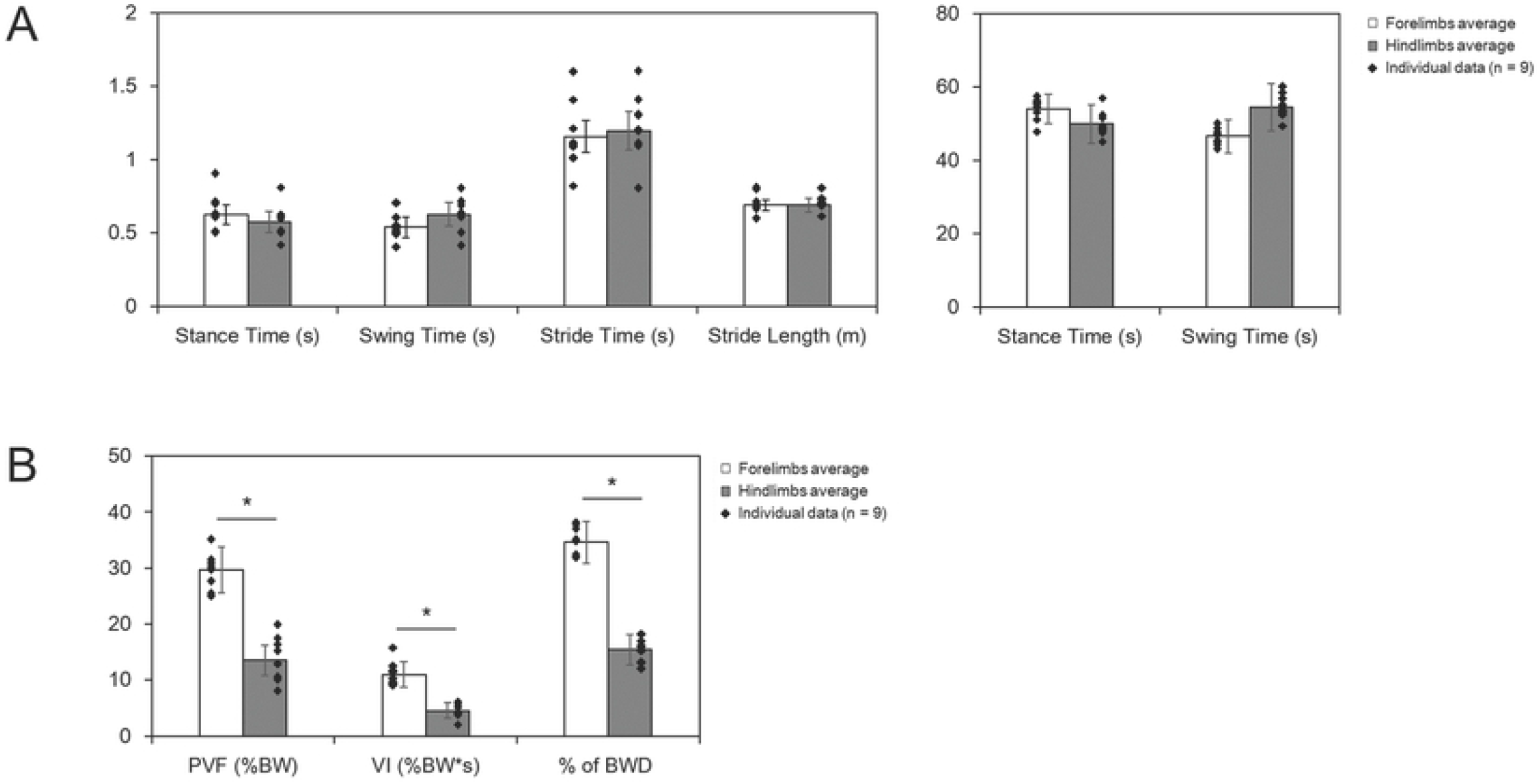
Comparison of the gait parameters for the forelimbs and hindlimbs. (A) Gait data for the temporo-spatial parameters (stance time, swing time, stride time, percentage of stance, percentage of swing, stride length). (B) Gait data for the kinetic parameters (PVF, peak vertical force; VI, vertical impulse; percentage of BWD, body weight distribution). Data are presented as mean ± SD, n = 9 in total. *p < 0.05 compared with forelimbs values (Kruskal-Wallis test).

### Comparison of gait parameter data for the forelimbs and hindlimbs

The averages of the forelimb and hindlimb gait parameters are presented in Fig 3. Scatter plots indicate individual data evaluated from each of the nine rhesus monkeys. Gait data for the temporo-spatial parameters show no differences between the forelimbs and hindlimbs (Fig 3A). However, most forelimb kinetic parameter values were generally higher than hindlimb kinetic parameter values (Fig 3B). Significant differences were observed in the PVF, VI, and percentage of BWD values.

### Assessment of gait symmetry

Tables 1 and 2 show the averages of the left- and right-side gait parameter values obtained from the nine monkeys. There were no significant differences between the temporo-spatial and kinetic data for the left and right forelimbs or the left and right hindlimbs (S1 Fig). The SI was used to evaluate the changes in gait symmetry. The median values of all symmetry indices were nearly zero, which indicates near perfect symmetric gait pattern (Table 3). There were no significant differences between the symmetry indices of the forelimbs and hindlimbs (Fig 4). These results suggest that most gait data obtained from the right and left side are similar for both forelimbs and hindlimbs of healthy rhesus monkeys.

**Fig 4.**
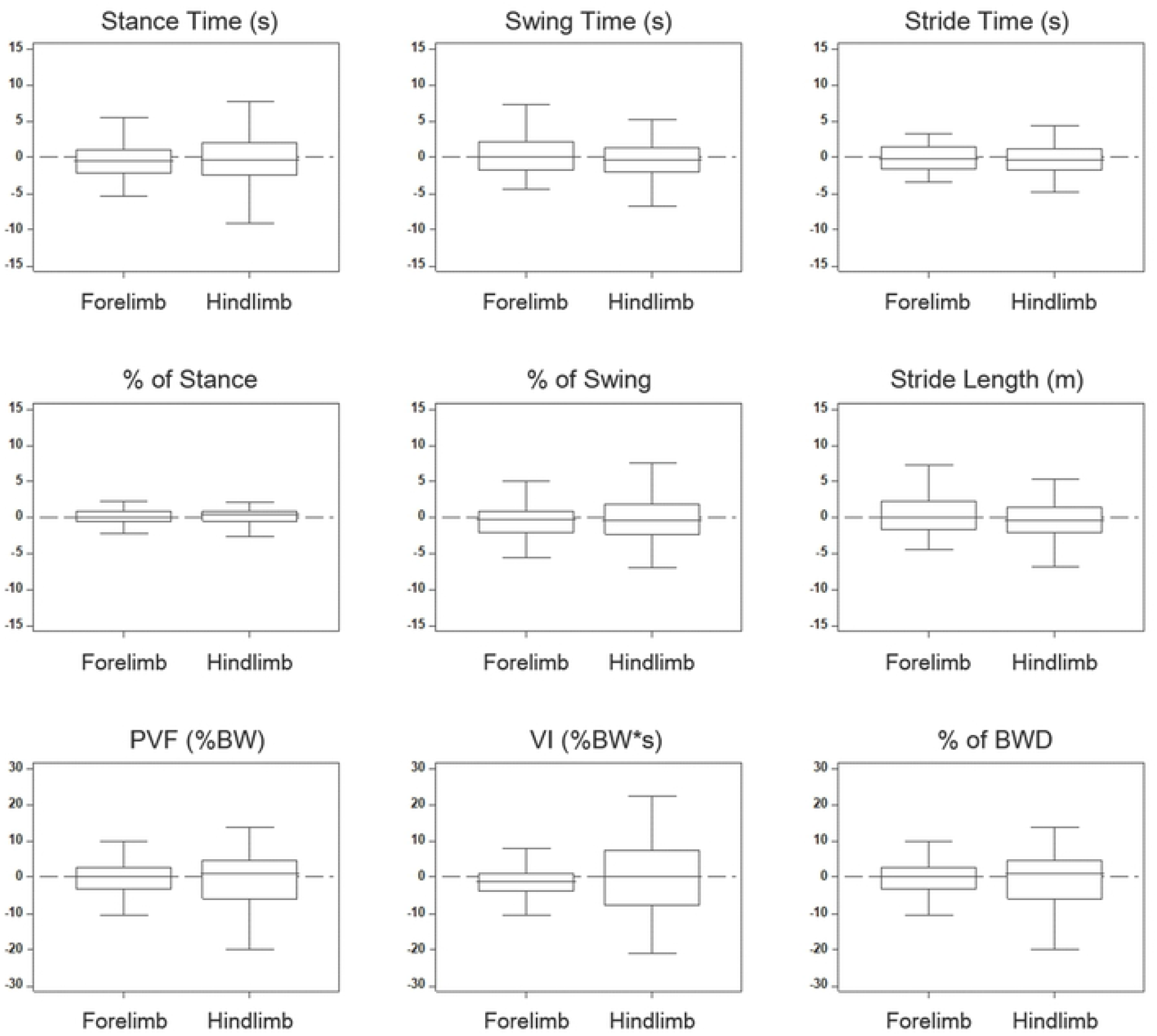
Box plots of gait data. The symmetry indices of gait parameters are visualized as box plots with median, interquartile range, and maximum and minimum values.

**Table 1.** Average gait data of left and right forelimbs obtained from nine rhesus monkeys

| Parameter |  | Left |  | Right |  |
| --- | --- | --- | --- | --- | --- |
| | | Mean ± SD | CV | Mean ± SD | CV |
| Temporo-spatial | Stance Time (s) | 0.63 ± 0.14 | 21.83 | 0.62 ± 0.11 | 17.28 |
| | Swing Time (s) | 0.54 ± 0.13 | 23.83 | 0.53 ± 0.11 | 20.09 |
| | Stride Time (s) | 1.17 ± 0.26 | 22.31 | 1.15 ± 0.21 | 18.65 |
| | Stride Length (m) | 0.68 ± 0.06 | 9.67 | 0.69 ± 0.05 | 8.58 |
| | % of Stance | 54.08 ± 5.03 | 9.31 | 54.06 ± 2.48 | 4.58 |
| | % of Swing | 46.72 ± 4.63 | 9.91 | 46.36 ± 3.22 | 6.95 |
| Kinetic | PVF (%BW) | 29.63 ± 2.69 | 9.08 | 29.69 ± 4.01 | 13.49 |
| | VI (%BW*s) | 11.03 ± 2.80 | 25.39 | 10.89 ± 2.71 | 24.91 |
| | % of BWD | 34.65 ± 2.26 | 6.51 | 34.50 ± 3.95 | 11.45 |

**Table 2.** Average gait data of left and right hindlimbs obtained from nine rhesus monkeys

| Parameter |  | Left |  | Right |  |
| --- | --- | --- | --- | --- | --- |
| | | Mean ± SD | CV | Mean ± SD | CV |
| Temporo-spatial | Stance Time (s) | 0.57 ± 0.10 | 18.36 | 0.58 ± 0.10 | 18.06 |
| | Swing Time (s) | 0.64 ± 0.13 | 20.45 | 0.61 ± 0.11 | 17.77 |
| | Stride Time (s) | 1.21 ± 0.24 | 19.56 | 1.18 ± 0.20 | 16.59 |
| | Stride Length (m) | 0.68 ± 0.06 | 9.46 | 0.69 ± 0.05 | 7.94 |
| | % of Stance | 49.34 ± 4.37 | 8.85 | 50.45 ± 3.25 | 6.44 |
| | % of Swing | 55.70 ± 4.66 | 8.36 | 53.23 ± 4.67 | 8.77 |
| Kinetic | PVF (%BW) | 13.05 ± 3.27 | 25.03 | 13.96 ± 3.85 | 27.58 |
| | VI (%BW*s) | 4.51 ± 1.19 | 26.51 | 4.58 ± 1.33 | 29.02 |
| | % of BWD | 15.00 ± 2.15 | 14.33 | 15.85 ± 2.81 | 17.70 |

**Table 3.**
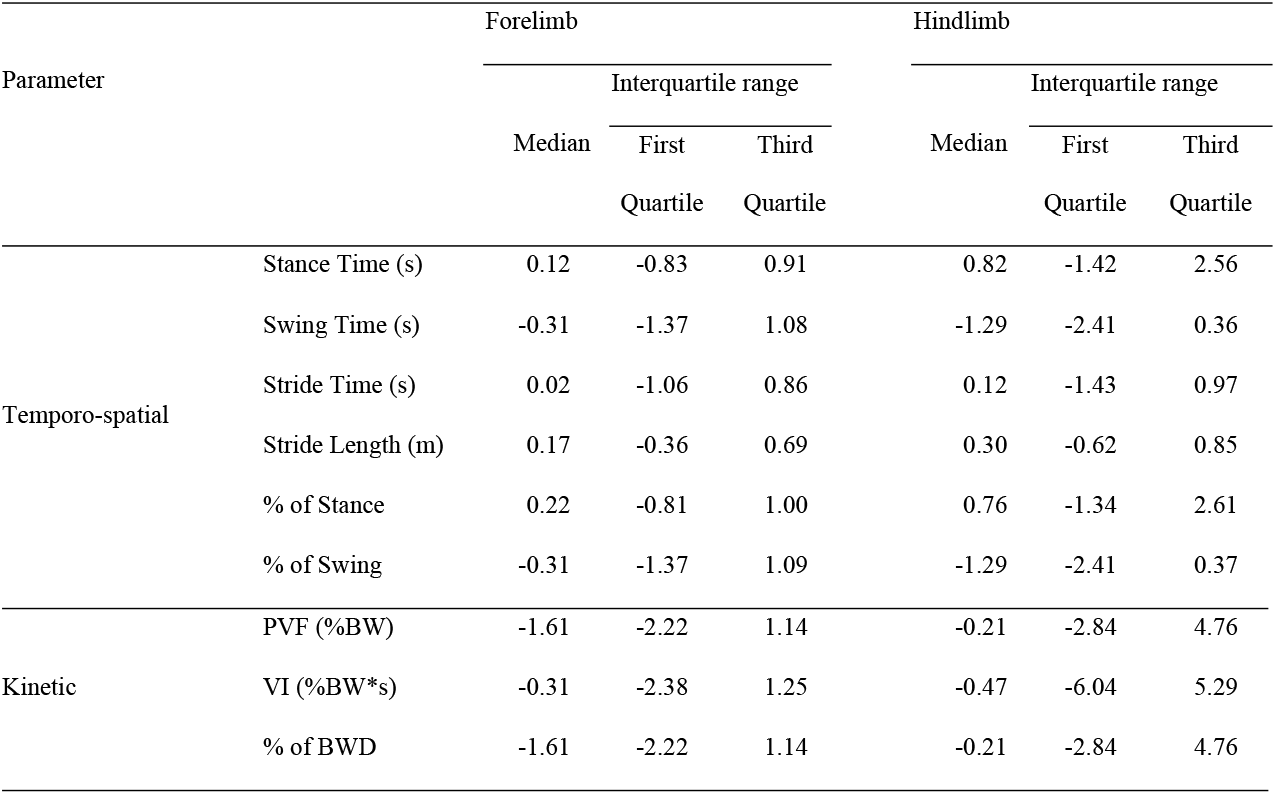
Comparison of the symmetry indices (%) of the temporo-spatial and kinetic parameters between the forelimbs and hindlimbs

## Discussion

Gait analysis in non-human primates with quadrupedal locomotion has been used to elucidate the role of the sensory-motor circuit in controlling locomotor movement and compare the species-specific locomotion with that of sub-primate quadrupedal animals [22]. Rhesus monkeys possess bipedal ability, although quadrupedal locomotion is used primarily [23]. In our experience too, the monkeys frequently demonstrated bipedal locomotion during gait analysis. A custom-built transparent plexiglass tunnel was used for stimulating habitual quadrupedal locomotion, and the gait parameters were measured while the animals walked in a comfortable environment.

This is the first study to evaluate and access temporo-spatial and kinetic data for rhesus monkeys using a pressure-sensing walkway system. This sensor system has been widely applied to the gait study of very small to large animals and has also used in clinical human studies, because it provides automated calculation of gait parameters without additional technical equipment, such as markers on the skin and calibrated camera setups [24–26]. Our results showed that there were no significant differences between temporo-spatial and kinetic gait parameter values, but there were detectable right-left symmetries between the limbs on the opposite side. As the SI is an objective parameter in normal gait function, the lack of asymmetry is a sign of a healthy rhesus gait, which might be used to detect dysfunction gait in such animals.

To avoid the variability in gait parameters, we controlled several variables, including type of locomotion, velocity, environmental condition, body weight, gender, and age. Studies using canine models show that walking speed could affect the ground reaction forces [27]. With an increase in gait velocity, the PVF increased and VI decreased [28]. The gait data from the present study showed low variability as determined by the inter-animal CVs.

The characteristics of the quadrupedal movement pattern while walking have been demonstrated for several different species of rodents, dogs, cats, sheep, and monkeys [21, 29–32]. In this study, our results showed that kinetic parameter values, including the PVF, VI, and percentage of BWD, are higher for the forelimbs than those for the hindlimbs. This pattern is similar to that reported for dogs, cats, and sheep. In principle, the weight difference between forelimbs and hindlimbs was observed during the animals walked down in a hall way. While walking, the body weight of sub-primate animals reportedly distributes approximately 30% to each forelimb and 20% to each hindlimb [21, 29, 31]. However, our results showed that the mean of percentage of BWD for forelimbs was higher in the present study than those in previous studies on other species [30, 33]. In this study, for rhesus monkeys walking at a speed of 0.62 m/s, the percentage of BWD for each forelimb and hindlimb was 35% and 15%, respectively. In contrast, the opposite pattern was observed while monkeys walked on a raised horizontal pole [34]. While walking on the pole, the reported PVF in monkeys was higher on hindlimbs than forelimbs, which indicates the influence of surface on the functional role of the four limbs.

One limitation of this study is that gait analysis was performed in single species (rhesus monkey) and on animals of a specific age (8 years). Generally, gait characteristics such as stride time, length, and velocity change with age in humans [35]. Furthermore, our sample size was quite small, and there was no study of pathological gait. Thus, further studies with larger sample sizes are necessary to validate our results and to investigate the differences in gait parameters between normal and unhealthy subjects.

## Conclusion

In this gait analysis conducted on healthy rhesus monkeys, there were no significant differences between the temporo-spatial and kinetic parameters. In addition, most SI values related to gait parameters showed a low variability, reflecting a symmetric gait pattern.

However, all forelimb kinetic parameters had high values. The results of our preliminary study obtained using a pressure-sensing walkway could be useful for distinguishing between normal and pathologic gait patterns in rhesus monkeys.

## Acknowledgements

The authors would like to thank Junghak Park at WinnerTech (Daejeon, Republic of Korea) for assistance with the custom-built transparent plexiglass tunnel. We also thank Kwon Yi at DooRee System Technology (Seongnam, Republic of Korea) for assistance with data collection.

## Supporting information

**S1 Table. Characteristics of experimental animals.**

**S2 Table. Temporo-spatial parameters of the forelimbs of individual rhesus monkeys.**

**S3 Table. Temporo-spatial parameters of the hindlimbs of individual rhesus monkeys.**

**S4 Table. Kinetic parameters of the forelimbs of individual rhesus monkeys.**

**S5 Table. Kinetic parameters of the hindlimbs of individual rhesus monkeys.**

**S1 Fig. Comparison of the gait parameter values of the forelimbs and hindlimbs for the left and right side.** Temporo-spatial data (stance time, swing time, stride time, percentage of stance, percentage of swing, stride length) and kinetic data (PVF, peak vertical force; VI, vertical impulse; percentage of BWD, body weight distribution) while walking represented as averaged values with standard deviations. *\*p* < 0.05 compared with forelimb values (Kruskal-Wallis test).

**S1 Video. A supplementary video of monkey gaits that pre-trained to walk on the pressure-sensing walkway positioned inside the custom-built tunnel.**

**S2 Video. Visualization of the performed steps using the pressure-sensing walkway system.** The provided software program automatically shows the placement of each of the four limbs.

